# MARS: a tool for haplotype-resolved population-based structural variation detection

**DOI:** 10.1101/2021.09.27.462061

**Authors:** Lu Zhang, Arend Sidow, Xin Zhou

**Affiliations:** Department of Computer Science, Hong Kong Baptist University, Kowloon Tong, Hong Kong; Department of Pathology, Stanford University, Stanford, CA 94305, USA; Department of Biomedical Engineering, Vanderbilt University, Nashville, TN 37235, USA; Department of Computer Science, Vanderbilt University, Nashville, TN 37235, USA

## Abstract

**Motivation:** Linked-reads enables genome-wide phased diploid assemblies. These haplotype-resolved assemblies allow us to genotype structural variants (SVs) with a high sensitivity and be able to further phase them. Yet, existing SV callers are designed for haploid genome assemblies only, and there is no tool to call SV from a large population of diploid assemblies which can define and refine SVs from a global view.

**Results:** We introduce MARS (Multiple Alignment-based Refinement of Svs) in linked-reads for the detection of the most common SV types - indels from diploid genome assemblies of a large population. We evaluated SVs from MARS based on Mendelian law of inheritance and PacBio HiFi reads and it achieved a high validation rate around 73%-87% for indels that we have selected from 34 assembled samples.

**Availability:** Source code and documentation are available on https://github.com/maiziex/MARS.

**Supplementary information:** Supplementary data are available at *Bioinformatics* online.

## 1 Introduction

The ultimate goal of diploid human genome sequencing is to obtain a comprehensive, and ideally 100% accurate, list of phased genetic variation harbored by each individual’s ‘germline’ genome (Levy *et al*., 2007). Genetic variation originates via a multitude of molecular mechanisms and takes many forms, including single nucleotide variants (SNPs) and the most common types of structural variants (SVs), insertions and deletions (indels). Linked-read sequencing has recently emerged as a potentially cost-effective solution for genome reconstruction and variation discovery. It is based on the idea that barcoded short reads can be generated from original long fragments of the DNA preparation such that all short reads from the same fragment have the same barcode. The approach was pioneered by 10x Genomics (Zheng *et al*., 2016). Even though their linked-reads has been discontinued recently, several companies have put effort into this promising technology, such as single tube long fragment reads (stLFR) by BGI (Wang *et al*., 2019), TELL-Seq by Universal Sequencing Technology (Chen *et al*., 2020), etc. Linked-read sequencing has low base pair-level error rates, and if the data are leveraged appropriately with custom algorithms, they facilitate discovery of SNPs and indels with good accuracy, as well as long-range haplotyping.

Regardless of technology employed, cost-effective and accurate indel discovery in personal genomes is still a challenge. While benchmarking standards have been developed for a handful of reference genomes, the confidence in each individual indel call is difficult to estimate (Chapman *et al*., 2020; Zook *et al*., 2020). An additional challenge is the precise and accurate determination of breakpoints, which is sometimes further complicated by compound events such as insertions that also delete short sequences around the insertion site (Cameron *et al*., 2019). This scenario could be pretty complicated and messy when calling SVs for a large population. A final complication is the fact that the molecular mechanism that gave rise to the indel allele is often the opposite of the call, whenever the reference genome carries the derived allele and the individual’s sequenced genome has the ancestral allele.

We here address some of these challenges by taking a multiple alignment-based approach to inform accuracy and precision of indel discovery through a tool called MARS. We performed reference-assisted whole genome diploid assembly of 34 linked-read genomes using Aquila, discovered indels (*≥* 20bp by default) in each assembled genome, and then assessed reliability of our calls on the basis of multiple sequence alignments of the indels and surrounding sequence.

**Fig. 1.**
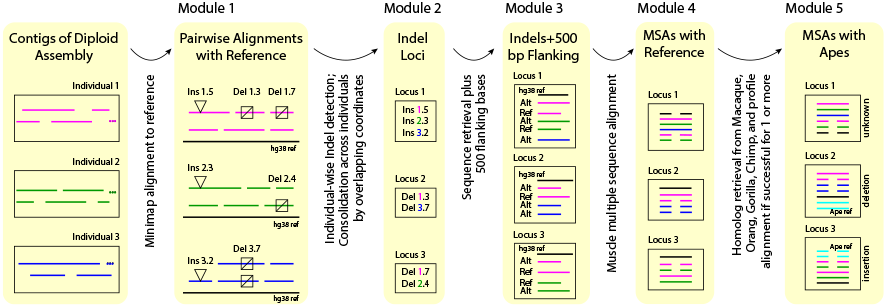
Schematic diagram of the last MSA module. The detailed descriptions are in the methods section.

## 2 Methods

MARS is a haplotype-resolved population-based structural variation detection method for linked-read diploid assemblies. The input files are whole genome diploid contigs from each sample. Aquila (Zhou *et al*., 2021) or Aquila_stLFR (Liu *et al*., 2021) can be used to assemble whole genome diploid contigs on 10x and stLFR linked-reads data, respectively. In this paper, we have assembled 34 samples of 10x linked-reads data with Aquila (see details in Supplementary Information).

The first module of MARS performs a pairwise contig-to-reference comparison to generate genome-wide variant calls from each haplotype independently. It then merges variants from both haplotypes to generate diploid variant calls. This step is performed for each sample. In the second module, MARS merges SVs of all samples through overlapping coordinates and a hard threshold (1bp for deletion and 100bp for insertion). The third module extracts haplotype-resolved sequences from the samples that contribute their original SVs to the merged SVs. To extract the haplotype-resolved sequences around the SV for each sample, it refers back to breakpoints of the original SV and extracts the SV sequence plus the left and right 500bp flanking regions from the assembled haplotype-resolved contigs. There are two scenarios to extract the haploid sequences for each sample. If the SV is homozygous or compound-heterozygous, MARs extracts both haploid sequences simply based on the breakpoints of the original SV from the sample. If the SV is heterozygous, it needs to extract the haploid sequence covering the reference allele and the haploid sequence covering the actual SV. For each merged SV, it also extracts the human reference sequence based on the breakpoints of the merged SV. Because the breakpoints of the new merged SV span the breakpoints of all original SVs, the reference sequence it extracts enclose all individual haploid sequences. In the fourth module, MARS uses MUSCLE (Edgar, 2004) to perform a multiple sequence alignment (MSA) of the reference sequence and all extracted haplotype-resolved sequences. It then scans through all MSAs to determine the outer breakpoints of each new merged SV from a global scale. To detect the breakpoints, it discovers the first large gap (*≥*20bp) on all sequences from left and right side, respectively. An optional fifth module of MARS facilitates investigation of the ancestral state of indels, by profile-aligning the homologous portions of Apes reference sequences (for example, Orangutan, Chimpanzee, Macaque or Gorilla) to each MSA.

## 3 Results and Discussion

We collected the linked-reads from publicly available 26 samples and sequenced 8 additional samples (Supplementary Note and Supplementary Table 1) to evaluate the performance of MARS. We selected SVs supported by at least two samples for later analyses. We adopted two strategies to validate SV calls: Mendelian inheritance and PacBio HiFi reads (Supplementary Note). For inheritance, the average validation rates were 84.48% and 78.92% for deletions and insertions (Supplementary Table 2) based on the four trios in our samples (Supplementary Table 3); for the four samples with PacBio HiFi reads, an average of 73.03% of deletions and 86.66% of insertions (Supplementary Table 4) were validated. The overall AUROC (Area under the ROC Curve) based on the SV confidence scores was 0.848 by applying ten-fold cross validation to the SVs validated by PacBio HiFi reads.

We also annotated the SVs functionally with Ensembl Variant Effect Predictor (McLaren *et al*., 2016) (Supplementary Figure 1-5) and RepeatMasker (Tarailo-Graovac and Chen, 2009) (Supplementary Figure 6). Coding SVs were rarely in repeat sequences and their lengths were commonly a multiple of three, suggesting the SV breakpoints refined by MARS were accurate. The performance of the SV confidence scores was a little bit different across the SVs located in the different functional regions (Supplementary Figure 7). We further selected derived deletions and insertions supported by different large numbers of haplotype-solved sequences; consistent with evolutionary expectation, the vast majority of full length Alu sequences were derived insertions.

To the best of our knowledge, MARS is the first tool to detect and refine SVs from a large population of diploid assemblies in linked-reads, and this approach could be further applied to long reads in the near further.

## Supporting information

MARS_SI

