## Supplementary material for "MARS: a tool for haplotype-resolved population-based structural variation detection": MARS_SI

### Supplementary Information

**Collect and sequence linked-read sequencing data**

We collected the publicly available 10x linked-read sequencing data of 26 samples from diverse populations, that were downloaded from 10x official data release (https://support.10xgenomics.com/de-novo-assembly/datasets), 1000 Genomes project (https://www.internationalgenome.org/) and Wong et al (Wong, et al., 2018) (Supplementary Table 1). All the library statistics were calculated by our in-house script. Seven additional libraries were prepared and sequenced as before (Zhang, et al., 2019). The sequencing data can be found in NCBI (PRJNA753653).

**Validation of SV calls by HiFi reads and violation of Mendelian law of inheritance**

The SVs were examined by two approaches: (i) We applied svviz2 (Spies, et al., 2015) to analyze PacBio HiFi reads from NA12878, NA24385, NA24143 and NA24149. svviz2 aligned and compared the HiFi reads to the reference sequence and the reconstructed alternative allele of candidate SVs. Genotypes 0/1 and 1/1 confirmed our SV calls; the validated SVs were defined as the ones if none of the genotypes from these four samples violate the genotype from MSA results of MARS. (ii) We designed three rules to examine the SVs that violate the Mendelian inheritance in the four trios (Supplementary Table 3) using the genotype from MSA of MARS and paftools (https://github.com/lh3/minimap2/tree/master/misc), respectively. The rules are

1. If the genotype of son/daughter is 0/0, none of the parents could be 1/1.
2. If the genotype of son/daughter is 1/0, at least of the parents should be 1/0 or 1/1.
3. If the genotype of son/daughter is 1/1, none of the parents could be 0/0.

We aligned all the contigs generated by Aquila (Zhou, et al., 2021) to the reference genome by minimap2 and identified the SVs and their genotypes by paftools.

**Annotation of SV sequences**

Deletions and insertions were annotated as Alu sequences if they were between 250 and 350bp long and could be uniquely aligned to the Alu consensus sequence from the UCSC Genome Browser. We used RepeatMasker (Tarailo-Graovac and Chen, 2009) to annotate tandem repeats. The other functional SVs were generated by Ensembl Variant Effect Predictor (VEP) (McLaren, et al., 2016).

### Supplementary Tables

| Sample | Valid Del | Validation Rate (Del) | Valid Ins | Validation Rate (INS) |
|---|---|---|---|---|
| **HG00514** | 25,919 | 81.92% | 15,652 | 78.81% |
| **HG00733** | 25,143 | 79.93% | 14,254 | 71.51% |
| **NA19240** | 29,764 | 88.81% | 19,236 | 82.06% |
| **NA24385** | 23,764 | 81.27% | 14,990 | 83.30% |

**Supplementary Table 2.** The validation rates of deletions and insertions based on Mendelian law of inheritance (the gap size of MSA should be larger than 20bp).

| Child | Father | Mother |
|---|---|---|
| HG00514 | HG00512 | HG00513 |
| HG00733 | HG00731 | HG00732 |
| NA19240 | NA19239 | NA19238 |
| NA24385 | NA24149 | NA24143 |

**Supplementary Table 3.** Four trios used to evaluate the performance of MARS.

| Samples | Valid Del | Validation Rate (Del) | Valid Ins | Validation Rate (Ins) |
|---|---|---|---|---|
| **NA24385** | 13,964 | 69.02% | 11,128 | 84.53% |
| **NA12878** | 14,230 | 74.23% | 12,423 | 89.47% |
| **NA24143** | 14,569 | 73.87% | 12,947 | 86.97% |
| **NA24149** | 14,398 | 74.99% | 12,869 | 85.66% |

**Supplementary Table 4.** The validation rates of deletions and insertions using PacBio HiFi reads (supported by at least two reads).

### Supplementary Figures

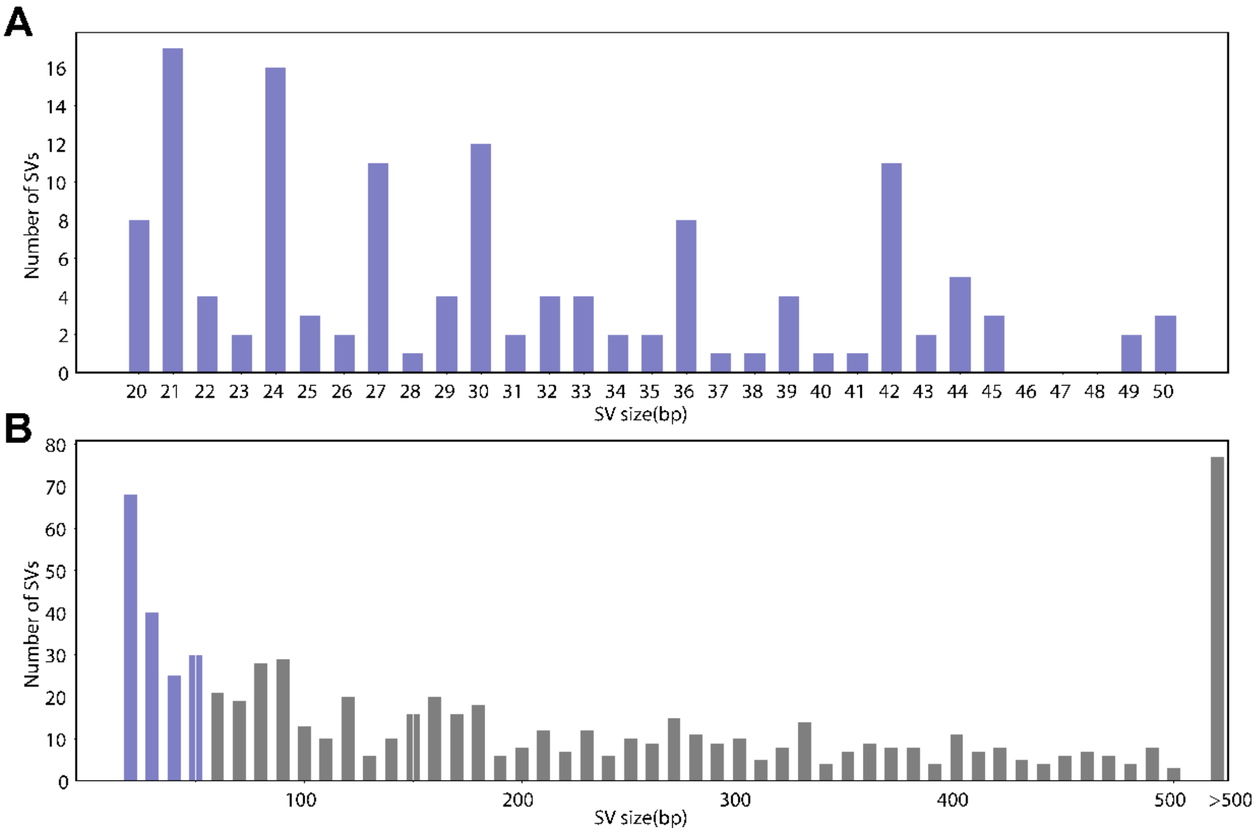


**Supplementary Figure 1.** The size distributions for the coding SVs between 20 to 50bp (**A**) and all coding SVs (**B**).

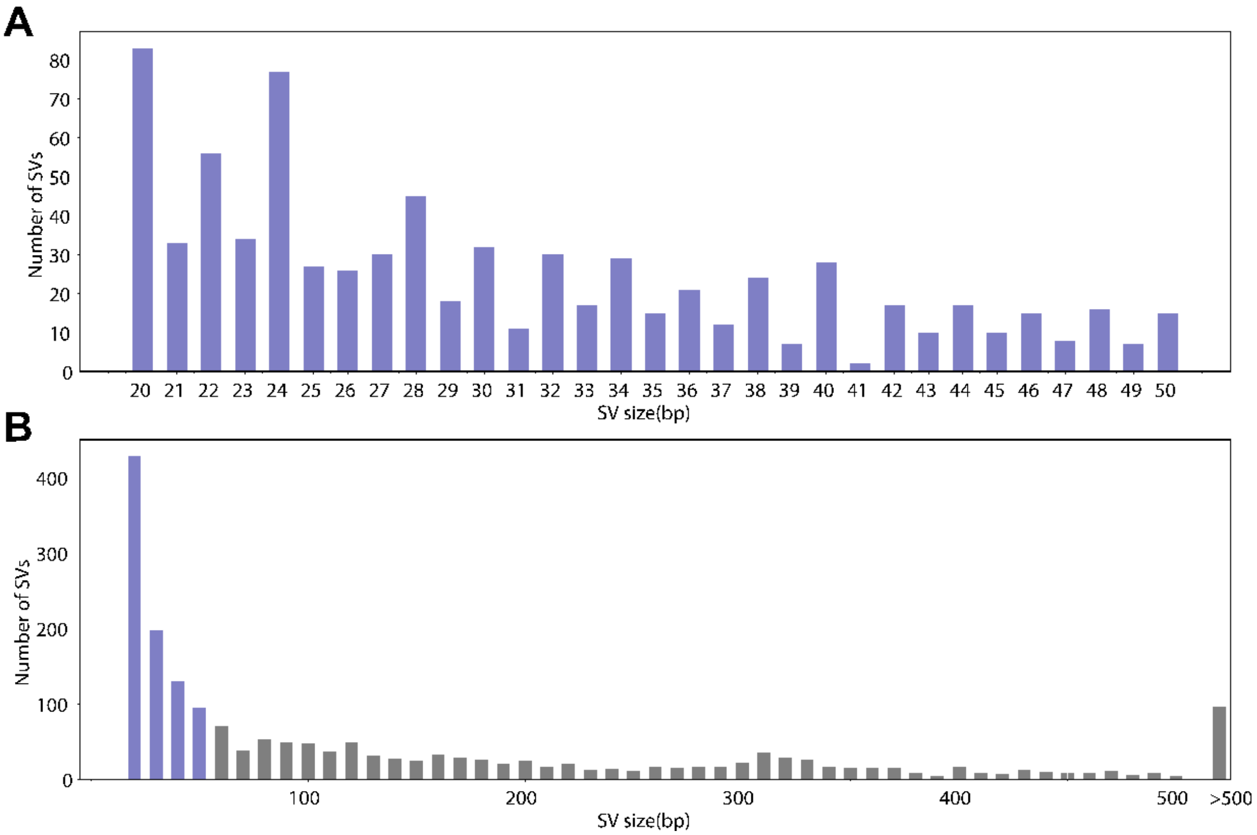


**Supplementary Figure 2.** The size distributions for the SVs from UTR between 20 to 50bp (**A**) and all the SVs from UTR (**B**).

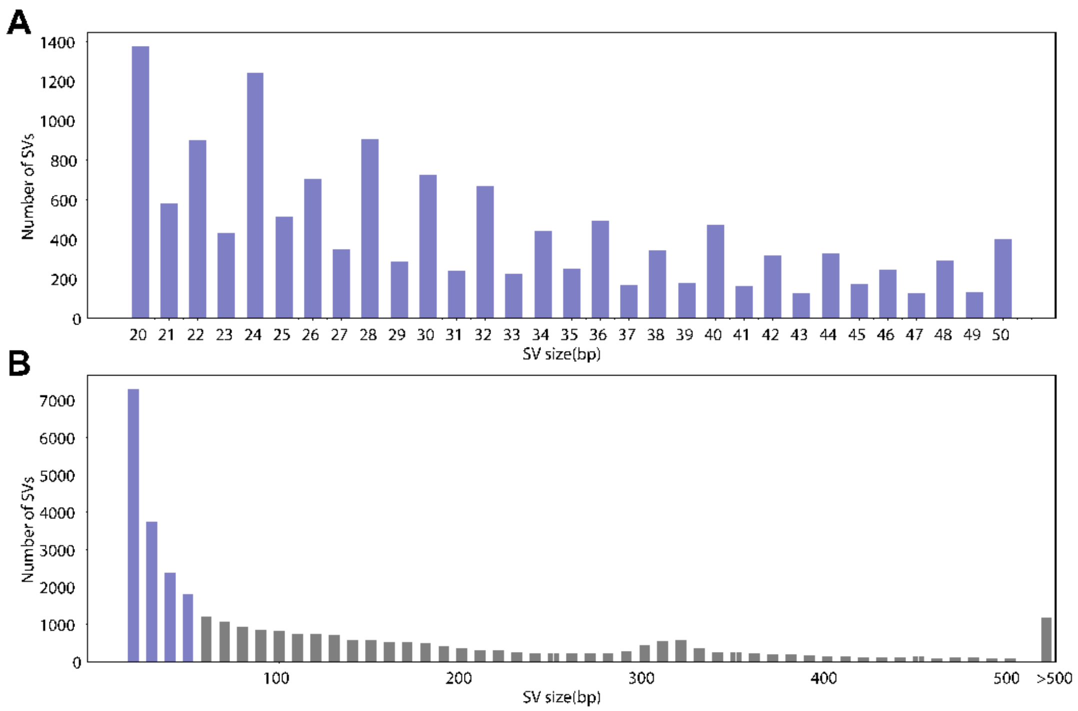


**Supplementary Figure 3.** The size distributions for the noncoding SVs from UTR between 20 to 50bp (**A**) and all the noncoding SVs (**B**).

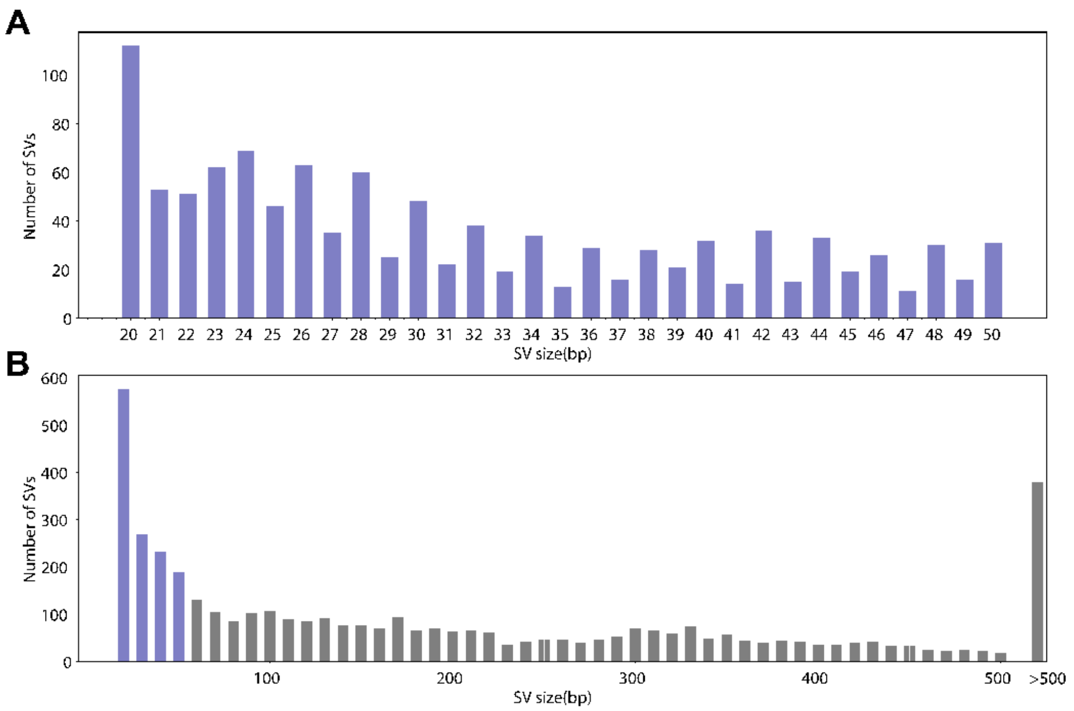


**Supplementary Figure 4.** The size distributions for the SVs from in transcription factor binding sites (TFBS) between 20 to 50bp (**A**) and all the SVs from in TFBS (**B**).

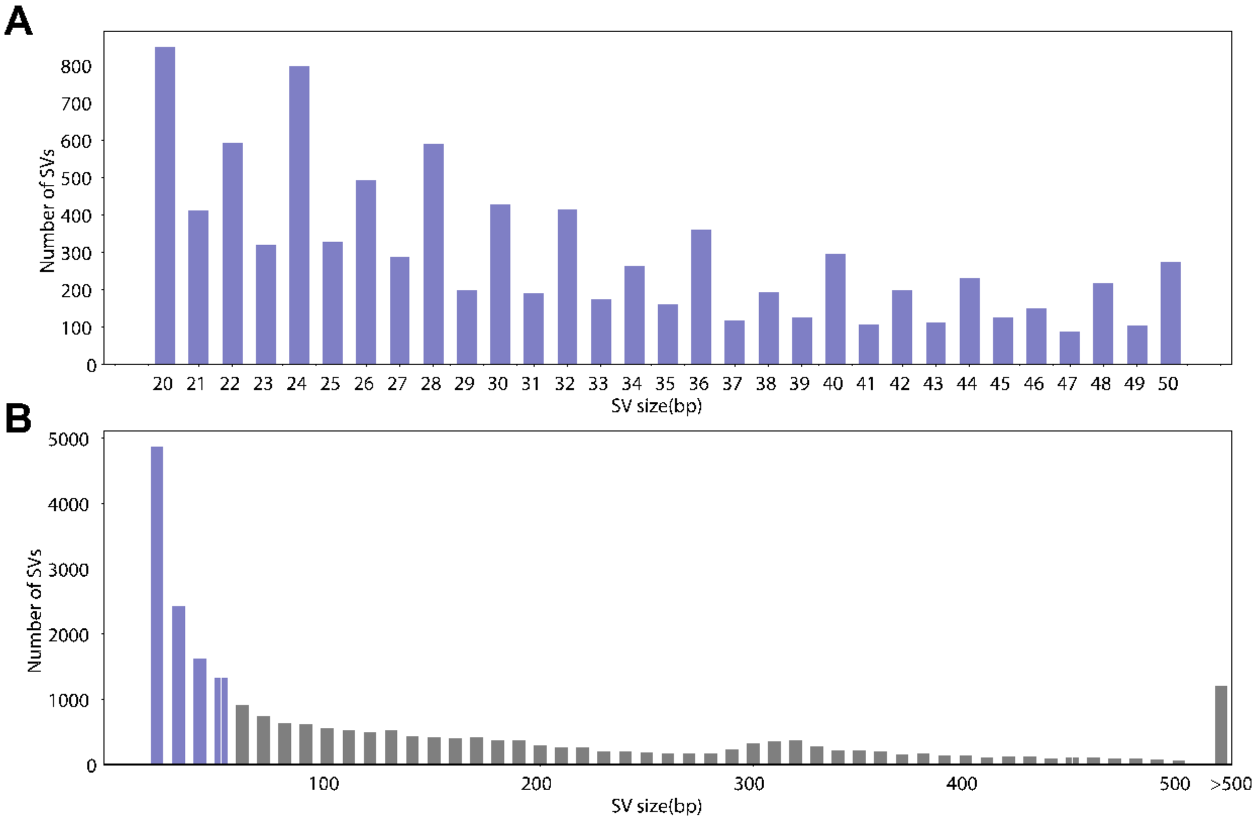


**Supplementary Figure 5.** The size distributions for the SVs from in regulatory regions between 20 to 50bp (**A**) and all the SVs from in regulatory regions (**B**).

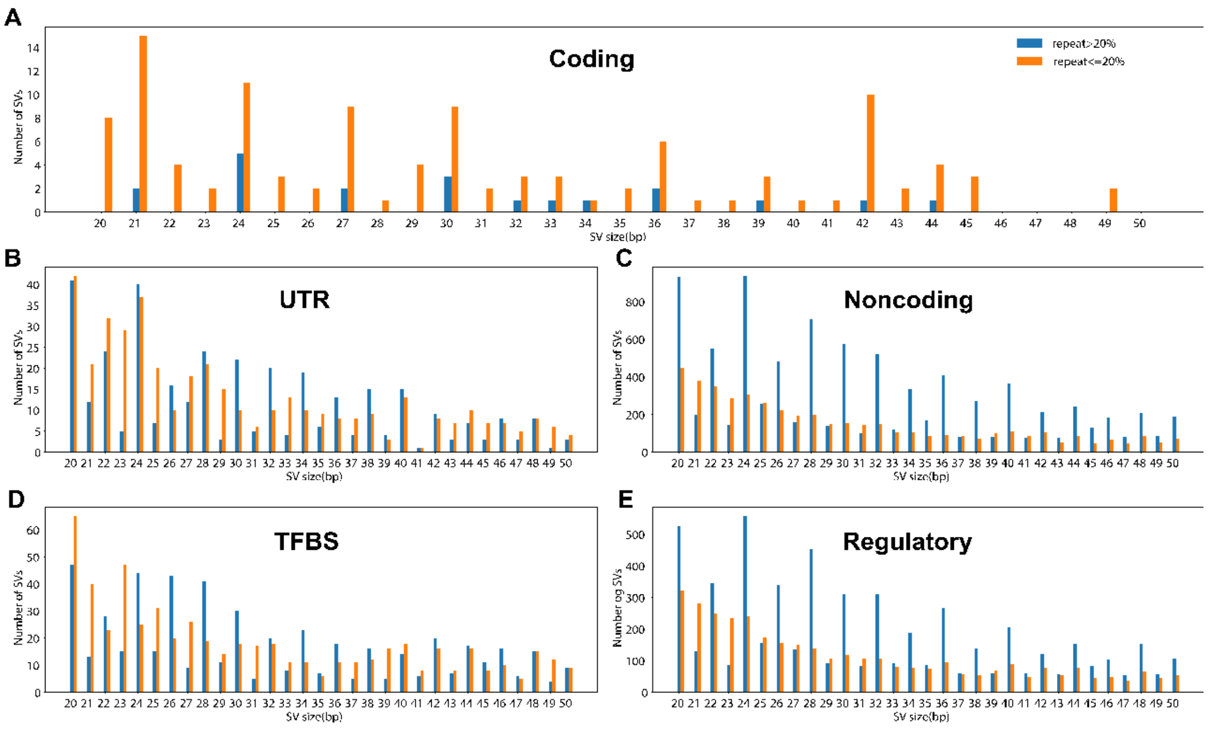


**Supplementary Figure 6.** Size distribution for (**A**) coding SVs, (**B**) SVs in UTR, (**C**) noncoding SVs, (**D**) SVs in transcription factor binding sites, (**E**) SVs regulatory regions (repeat>20% vs. repeat<=20%).

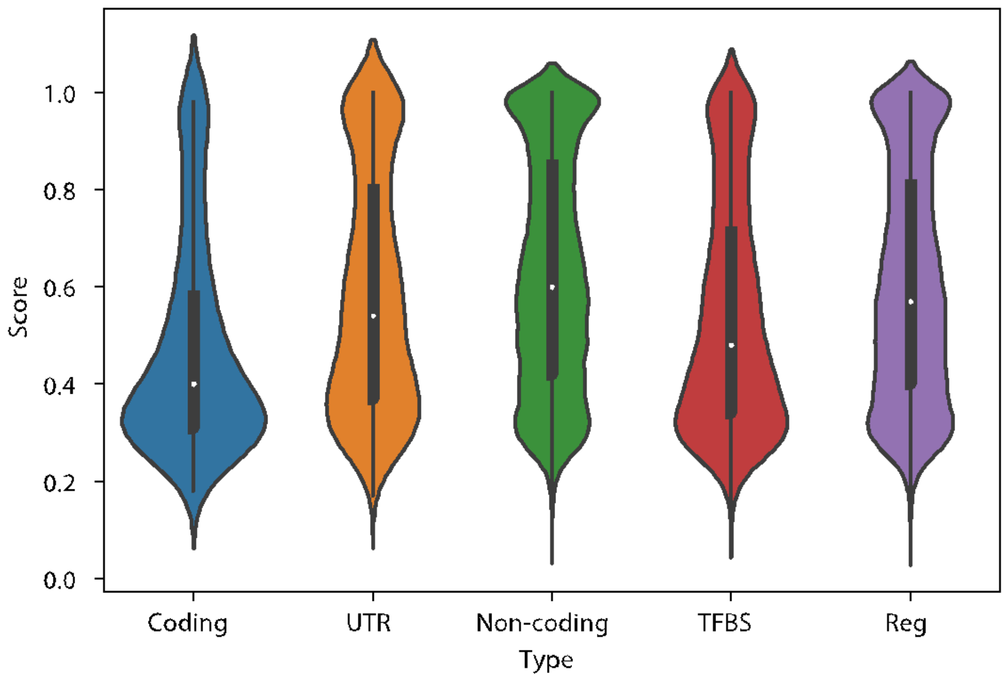


**Supplementary Figure 7.** Evaluation score distributions for the SVs from five categories.

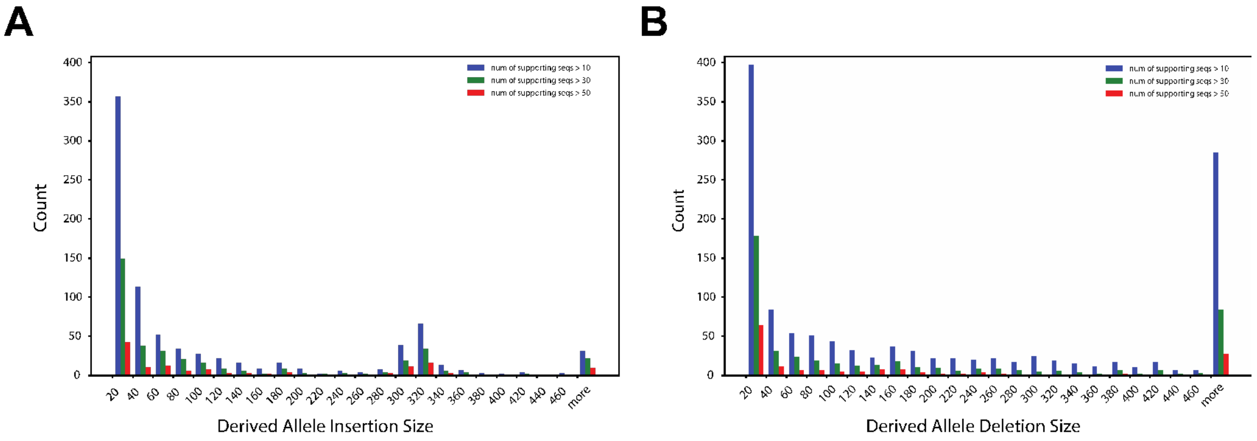


**Supplementary Figure 8**. (A) Derived allele insertion size distribution when number of supporting sequences is larger than 10 (blue bar), 30 (green bar) and 50 (red bar). (B) Derived allele deletion size distribution when number of supporting sequences is larger than 10 (blue bar), 30 (green bar) and 50 (red bar).
